# VariantStore: A Large-Scale Genomic Variant Search Index

**DOI:** 10.1101/2019.12.24.888297

**Authors:** Prashant Pandey, Yinjie Gao, Carl Kingsford

## Abstract

The ability to efficiently query genomic variants from thousands of samples is critical to achieving the full potential of many medical and scientific applications such as personalized medicine. Performing variant queries based on coordinates in the reference or sample sequences is at the core of these applications. Efficiently supporting variant queries across thousands of samples is computationally challenging. Most solutions only support queries based on the reference coordinates and the ones that support queries based on coordinates across multiple samples do not scale to data containing more than a few thousand samples. We present VariantStore, a system for efficiently indexing and querying genomic variants and their sequences in either the reference or sample-specific coordinate systems. We show the scalability of VariantStore by indexing genomic variants from the TCGA-BRCA project containing 8640 samples and 5M variants in 4 Hrs and the 1000 genomes project containing 2500 samples and 924M variants in 3 Hrs. Querying for variants in a gene takes between 0.002 – 3 seconds using memory only 10% of the size of the full representation.

---

Advanced sequencing technology and computing resources have led to large-scale genomic sequencing efforts producing genomic variation data from thousands of samples, such as the 1000 Genomes project [1–3], GTEx [4], and The Cancer Genome Atlas (TCGA) [5]. Analysis of genomic variants combined with phenotypic information of samples promises to improve applications such as personalized medicine, population-level disease analysis, and cancer remission rate prediction. Although numerous studies [6–12] have been performed over the past decade involving genomic variation, the ability to scale these studies to large-scale data available today and in the near future is still limited.

On an individual sample, the typical result of sequencing, alignment, and variant calling is a collection of millions of sample-specific variants. A variant is identified by the position in the chromosome where it occurs, an alternative sequence, a list of samples that contain the variant, and phasing information. The standard file format to report these variants is the variant call file (VCF) [13].

A common task is to identify all the samples with a given pattern of variants or identify all samples that have variants in a given gene. These tasks are translated into variant queries that require finding variants, samples with variants, or sample sequences between two positions in a chromosome. Applications often compare variants across multiple samples and need to perform the above queries based on sample-specific coordinate systems. A coordinate system is a system that uniquely identifies the positions of variants in a given genome. Each sample in variation data has a coordinate system that can be different from the reference coordinate system. Variants can appear at different positions in a sample coordinate system compared to the reference due to insertions and deletions (indels).

To effectively use variant information from many samples, medical and scientific applications must often answer many instances of one or more of the following types of queries:

1. Find the closest variant to position *X* for all samples in the reference coordinates.
2. Find the sequence between positions *X* and *Y* for sample *S* in the reference coordinates.
3. Find the sequence between positions *X* and *Y* for sample *S* in the sample coordinates.
4. Find all variants between positions *X* and *Y* for sample *S* in the reference coordinates.
5. Find all variants between positions *X* and *Y* for sample *S* in the sample coordinates.
6. Find all variants between positions *X* and *Y* for all samples in the reference coordinates.

These queries can be further used as building blocks for more complicated queries, such as finding samples with more than a given number of variants between two positions or count the number of variants for each sample between two positions.

Supporting variant queries based on multiple sample coordinate systems requires maintaining a function per sample that can map a position in the reference coordinate (as present in VCF files) to the sample coordinate.

Maintaining thousands of such functions requires storing and accessing an order of magnitude more data than only indexing variants based on a single reference coordinate system. Efficiently supporting thousands of coordinate systems adversely affects the memory footprint and computational complexity of the system making this problem much more challenging. This limits the scalability of variant indexes that support multiple coordinate systems to variation data containing only a few thousands samples.

VG toolkit [14] is one of the most widely used tools to represent genomic variation data and it also supports multiple coordinate systems. It encodes genomic variants from multiple samples in a graph, called a variation graph. A variation graph is a sequence graph where each node represents a sequence and a set of nodes through the graph, known as a path, embeds the complete sequence corresponding to the reference or a sample. Each node on a path is assigned a position indicating the location of the sequence in the coordinate system of the path. A node can be assigned multiple positions based on the number of paths that pass through the node. The variation graph enables read alignment against multiple sample sequences containing variants simultaneously and avoids mapping biases that arise when mapping reads to a single reference sequence [14–18].

VG toolkit stores each sample path as a list of nodes in the graph and maintains a separate index corresponding to the coordinates of the reference and samples. Storing a separate list of nodes for each sequence impedes the scalability of the representation for storing variation from thousands of samples. Moreover, variants are often shared among samples, so storing a list of nodes for each sample path introduces redundancy in the representation. VG toolkit is designed to optimize read alignment and uses sequence-based indexes for alignment. It can not be directly used for variant queries that require an index based on the position of variants in multiple sequence coordinates. Finally, the VG toolkit representation does not store phasing information contained in VCF files, which is required in many analyses.

Multiple solutions have been proposed that efficiently index variants and support a subset of the variant queries described above. GQT [19] was the first tool that proposed a sample-centric index for storing and querying variants. It stores variants in compressed bitmap indexes and supports efficient comparisons of sample genotypes across many variant positions. BGT [20] and GTC [21] proposed variant-centric indexes that store variants in compressed bit matrices. They support queries for variants in a given region and allow filtering returned variants based on subsets of samples. The SeqArray library [22] is another variant-centric tool for the R programming language to store and query variants.

However, these tools index variants only in the reference coordinate system and do not support variant queries in a sample coordinate system. Supporting multiple coordinate systems is a much harder problem to tackle. Furthermore, these tools do not store the reference sequence and cannot be directly used to query and compare sample sequences in a given region. Other tools have been proposed that use traditional database solutions, such as SQL and NoSQL [23–25]. However, they have proven prohibitively slow to index and query collections of variants.

We present VariantStore, a system for efficiently indexing and querying genomic information (genomic variants and phasing information) from thousands of samples containing millions of variants. VariantStore supports querying variants occurring between two positions across a chromosome based on the reference or a sample coordinate. VariantStore bridges the gap between the tools that are space-efficient and fast but only support reference-based queries (e.g., GTC [21]) and VG, which maintains multiple coordinate systems by storing variants in a variation graph but fails to scale to thousands of samples. We show this by indexing variants from both the 1000 genomes and TCGA projects (*>* 8K samples), and show that VariantStore is faster than VG and takes less memory and disk space. VariantStore performs variant queries based on sample coordinates in less than a second. Furthermore, we have designed VariantStore to efficiently scale out of RAM to storage devices in order to cater to the ever increasing sizes of available variation data by performing memory-efficient construction and query.

We encode genomic variation in a directed, acyclic variation graph and build a position index (a mapping of node positions to node identifiers) on the graph to quickly access a node in the graph corresponding to a queried position. Each node in the variation graph corresponds to a variant and stores a list of samples that contain the variant along with the position of the variant in the coordinate of those samples. The inverted index design allows one to quickly find all the samples and positions in sample coordinates corresponding to a variant. It also avoids redundancy that otherwise arises in maintaining individual variant indexes for each sample coordinate and scales well in practice when the number of samples grows beyond a few thousand.

To perform index construction and query efficiently in terms of memory, we partition the variation graph into small chunks (usually a few MBs in size) based on the reference coordinates and store variation graph nodes in these chunks. During construction, an active chunk is always maintained in memory in which new nodes are added, and once it reaches its capacity we compress and serialize it to disk and create a new active chunk. The nodes in and across chunks are ordered based on the reference coordinate since they are created based on variants in the VCF file which are themselves ordered by the reference coordinate. During a query, we only load the chunks in memory that contain the nodes corresponding to the query range.

To efficiently scale to thousands of coordinate systems (or samples), we maintain the position index only on the reference coordinate. The position index maps positions in the reference sequence where there is a variant to nodes corresponding to those variants in the variation graph. To lookup a position using a sample’s coordinate system, we first lookup the node corresponding to the position in the reference coordinate. We then traverse the sample path from the node in the graph to determine the appropriate node in the sample coordinate. A node with a given position in a sample coordinate is often close to the node in the reference coordinate with the same position.

## Results

Our evaluation of VariantStore is based on four parameters: construction time, query throughput, disk space, and peak memory usage.

To calibrate our performance numbers, we compare VariantStore against VG toolkit [26]. VG toolkit represents variants in a variation graph and supports multiple coordinate systems but does not support variant queries. Therefore, we only compare the construction performance and disk space against VG toolkit.

### Data

We use 1000 Genomes Phase 3 data [27] and three of the biggest projects from TCGA in terms of the number of samples, Ovarian Cancer (OV), Lung Adenocarcinoma (LUAD), and Breast Invasive Carcinoma (BRCA), for our evaluation. 1000 Genomes data contains more variants compared to the TCGA data but TCGA data contains more samples. 1000 Genomes data contains a separate VCF file for each chromosome containing variants from thousands of samples. The number of samples in each file is ≈ 2.5K. The variants in 1000 Genomes project are based on GRCh37 reference genome. The TCGA data contains a separate VCF file for each sample. The OV, LUAD, and BRCA projects contain 2436, 2680, and 4319 VCF files containing both normal and tumor variants respectively. For each project in TCGA, we first merged VCF files using the BCF tool merge command [28] and created a separate VCF file for each chromosome. The variants in the TCGA project are based on GRCh38 reference genome.

### Index construction

The total time taken to construct the variation graph representation and index includes the time taken to read and parse variants from the VCF file, construct the variation graph representation and indexes, and serialize the final representation to disk. For VariantStore, the reported time includes the time to create and serialize the position index. The space reported for VariantStore is the sum of the space of the variation graph representation and position index.

For VG toolkit, creating a variation graph representation with multiple coordinate systems (or sample path annotations) requires creating two indexes, XG and GBWT index. The XG index is a succinct representation of the variation graph without path annotations that allows memory- and time-efficient random access operations on large graphs. The GBWT (graph BWT) is a substring index for storing sample paths in the variation graph. We first create the variation graph representation using the “construct” command including all sample path annotations. We then create the XG index and GBWT index from the variation graph representation to create an index with all sample path annotations in the variation graph. For VG toolkit, the reported time includes the time to create and serialize the XG and GBWT indexes. The space reported for VG toolkit is the sum of the space of the XG and GBWT indexes. VG toolkit could not build GBWT index on TCGA data even after running for more than a day. We only report space for the XG index (which does not contain any sample path annotations) for TCGA data.

For both VariantStore and VG toolkit, we created 24 separate indexes, one each for chromosomes 1 *−* 22, and X and Y. Each of these indexes were constructed in parallel as a separate process. We report the time taken for construction as the time taken by the process that finishes last. For disk space, we report the total space taken by all 24 indexes. For peak memory usage, we report the highest individual and aggregate peak RAM usage for all processes.

The performance of VariantStore and VG toolkit for constructing the index on the 1000 Genomes and TCGA data is shown in Table 1.

**Table 1:** Time, space, peak RAM, and peak RAM (aggregate) to construct variant index on the 1000 Genomes and TCGA (OV, LUAD, and BRCA) data using VariantStore and VG toolkit. ^∗^Space for the XG index that does not contain any path information. We constructed all 24 chromosomes (1 – 22 and X and Y) in parallel. The time and peak RAM reported is for the biggest chromosome (usually chromosome 1 or 2). The space reported is the total space on disk for all 24 chromosomes. The peak RAM (aggregate) is the aggregate peak RAM for all 24 processes.

| System | Time | Disk space | Peak RAM | Peak RAM Agg. |
| --- | --- | --- | --- | --- |
| Dataset | 1000 Genomes |  |  |  |
| VariantStore | 3 Hrs 25 mins | 41 GB | 8.8 GB | 153 GB |
| VG-toolkit | 11 Hrs 10 mins | 50 GB | 37 GB | 450 GB |
| Dataset | TCGA (OV) |  |  |  |
| VariantStore | 1 Hr 5 mins | 3.4 GB | 1.1 GB | 17.45 GB |
| VG-toolkit |  | 11 GB* |  |  |
| Dataset | TCGA (LUAD) |  |  |  |
| VariantStore | 1 Hr 20 mins | 3.5 GB | 2.3 GB | 36.05 GB |
| VG-toolkit |  | 12 GB* |  |  |
| Dataset | TCGA (BRCA) |  |  |  |
| VariantStore | 4 Hrs 36 mins | 4.2 GB | 3.2 GB | 53.21 GB |
| VG-toolkit |  | 14 GB* |  |  |

VariantStore is 3× faster, takes 25% less disk space, and 3× less peak RAM than VG toolkit. For the TCGA data, VG toolkit could not build GBWT index embedding all sample paths. However, even the space needed for the XG index (not embedding sample paths) is ≈ 3.3× larger than the VariantStore representation containing all sample paths.

### Query throughput

We measured the query throughput for all six queries mentioned above. To show the robustness of query efficiency on different data we evaluate on two different chromosome indexes from two different projects. We chose Chromosome 2 which is one of the bigger chromosomes and Chromosome 22 which is one of the smaller ones. We evaluate query time on chromosome indexes from 1000 Genomes and TCGA LUAD data.

To perform queries, we specify a pair of positions in the reference or a sample coordinate system depending on the query type and optionally a sample name. Query parameters, such as the length of the queried region and density of variants in that region, affect the query timing. Therefore, we performed three sets of query benchmarks containing 10, 100, and 1000 queries and report the aggregate time. For each query in the set, we uniformly randomly pick the start position across the full chromosome length. The size of the query range is set to ≈ 42K bases which is approximately the length of a typical gene. Picking multiple query regions uniformly randomly across the chromosome provides a good coverage of different regions across the chromosome.

To measure the query throughput, we only load the position index and graph topology in memory and the variation graph representation is kept on disk before performing the query. Keeping the variation graph representation on disk keeps the peak memory usage low and all disk accesses are performed during the query to load appropriate node chunks.

The query throughput on 1000 Genomes data is shown in Figures 1a and 2a. For all query types, the aggregate time taken to execute queries increases linearly with the number of queries. Finding the sequence corresponding to a sample in a region (Queries #2, #3) takes less time compared to finding variants in a region (Queries #4, #5, #6). Finding the sequence takes less time because it involves traversing the sequence specific path in the region and reconstructing the sequence. However, finding variants in a given region takes more time because it involves an exhaustive search of neighbors at each node in the region to determine all the variants that are contained by a given sample or all samples.

**Figure 1:**
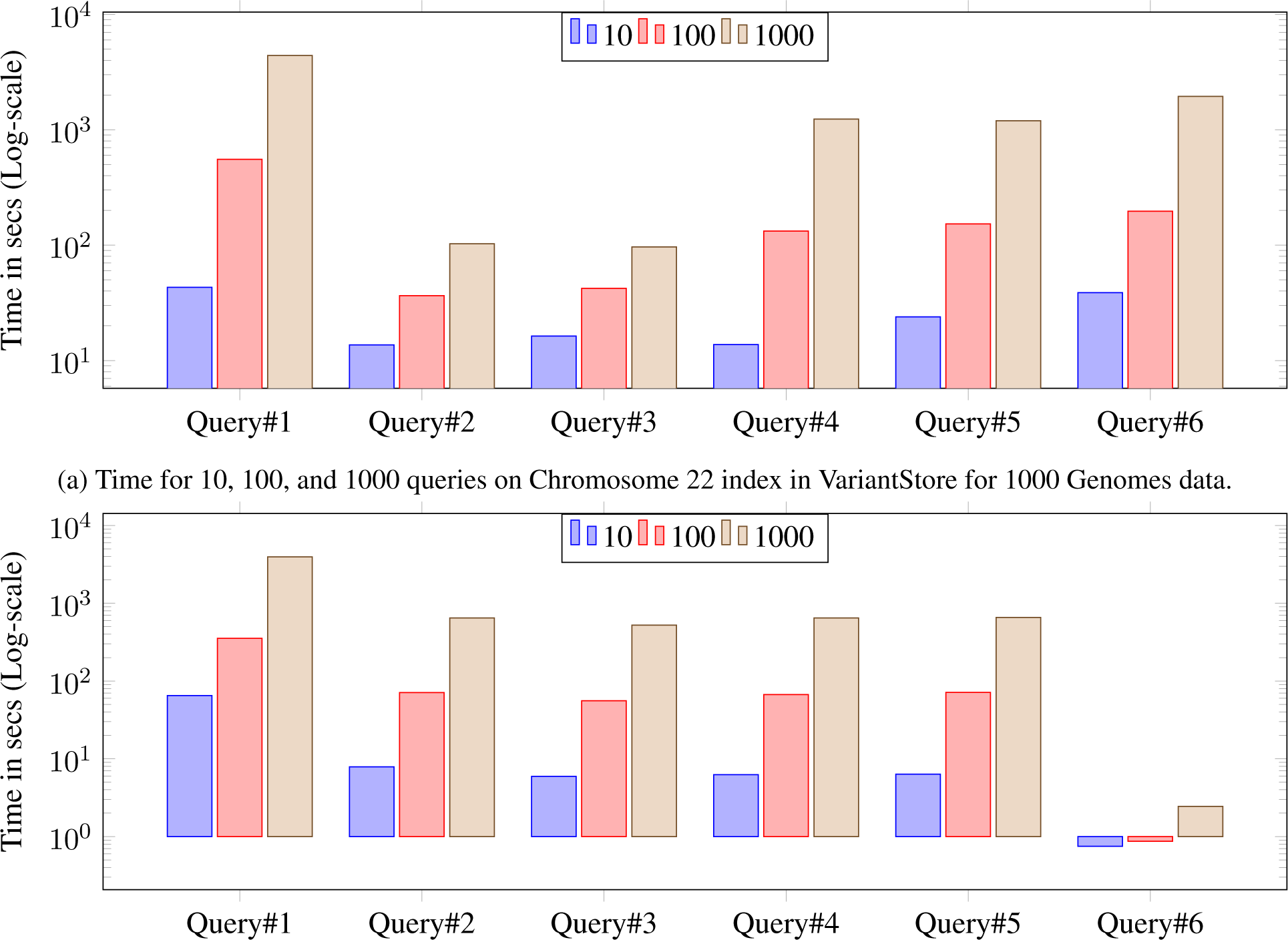
Time reported is the total time taken to execute 10, 100, and 1000 queries. For all queries the query length is fixed to ≈ 42*K*. Query#1: Closest variant, Query#2: Seq in ref coordinate, Query#3: Seq in Sample coordinate, Query#4: Sample variants in ref coordinate, Query#5: Sample variants in Sample coordinate, Query#6: All variants in ref coordinate

**Figure 2:**
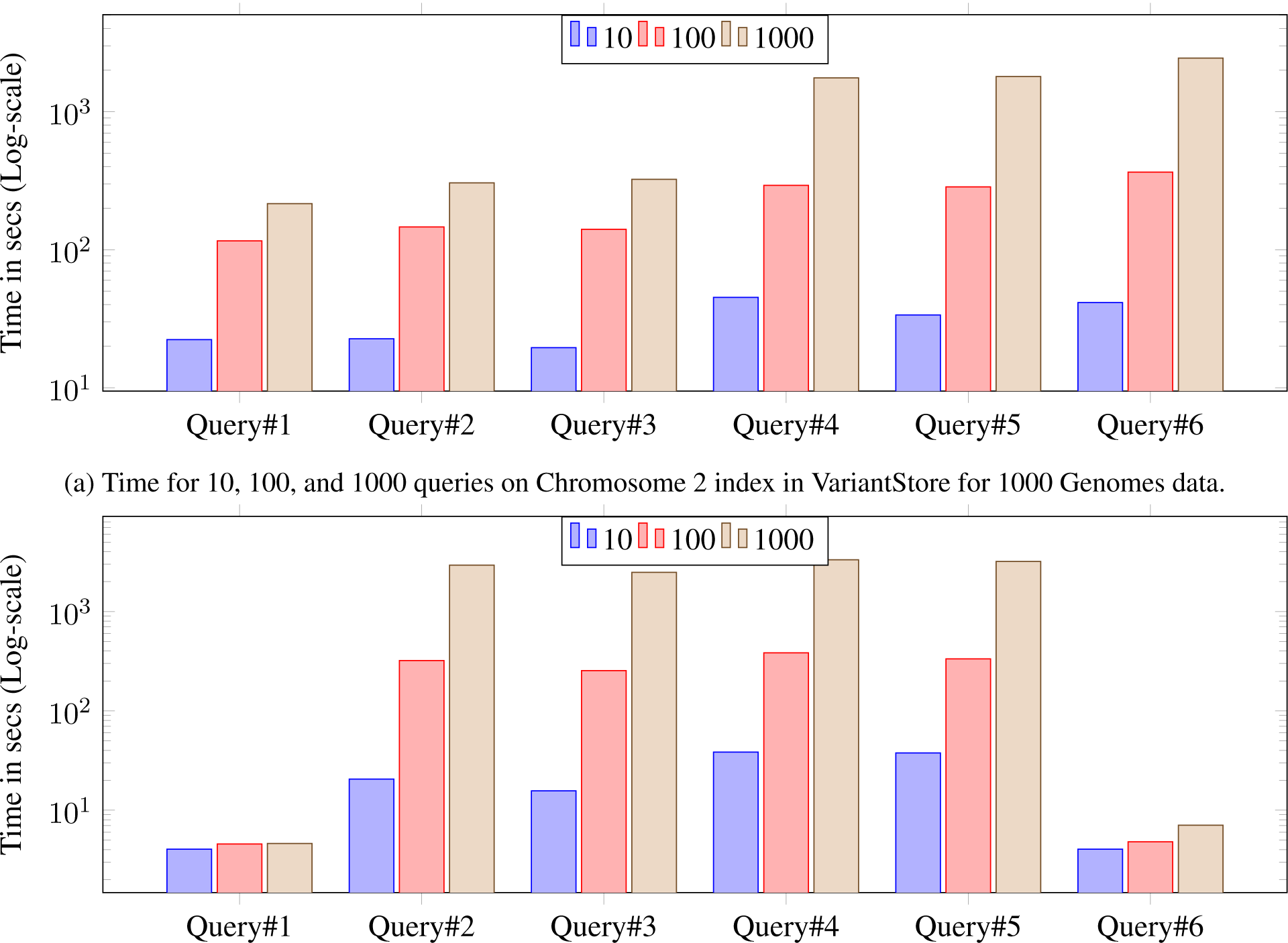
Time reported is the total time taken to execute 10, 100, and 1000 queries. For all queries the query length is fixed to ≈ 42*K*. Query#1: Closest variant, Query#2: Seq in ref coordinate, Query#3: Seq in Sample coordinate, Query#4: Sample variants in ref coordinate, Query#5: Sample variants in Sample coordinate, Query#6: All variants in ref coordinate

The query throughput on TCGA data is shown in Figures 1b and 2b. The TCGA data has twice as many samples compared to 1000 Genomes data but there are fewer variants. This makes the variation graph much sparser and queries in the position index become more expensive compared to traversing the graph between two positions. Finding all variants is the fastest query because the variation graph is very sparse and most position ranges are empty. Traversing the graph to find the sequence or variants for a sample takes similar amount of time. Finding the closest variant from a position takes the most amount of time because it involves performing multiple position-index queries to determine the closest variant.

For both 1000 Genomes and TCGA LUAD data, finding the closest variant query is faster for Chromosome 2 because variants in Chromosome 22 are more dense compared to Chromosome 2 which makes it faster to locate the closest variant.

### The effect of query range on peak memory usage

We performed another query benchmark to evaluate the effect of size of the position range on the peak memory usage and time. For this benchmark, we chose the “Sample sequence in reference coordinate” query (query #2) because this query involves traversing the full sample path between two positions. We performed sets of 100 queries with increasing size of the position range and record the total time and peak RAM usage. For each query in the set, we uniformly randomly pick the start position across the full chromosome length.

Effect of the query range size on peak memory usage and time is shown in Figure 3. The memory usage remains constant regardless of the query length. This is because during a query we access node chunks in sequential order and regardless of the query length only load at most two node chunks in RAM at a time. This keeps the memory usage essentially constant.

**Figure 3:**
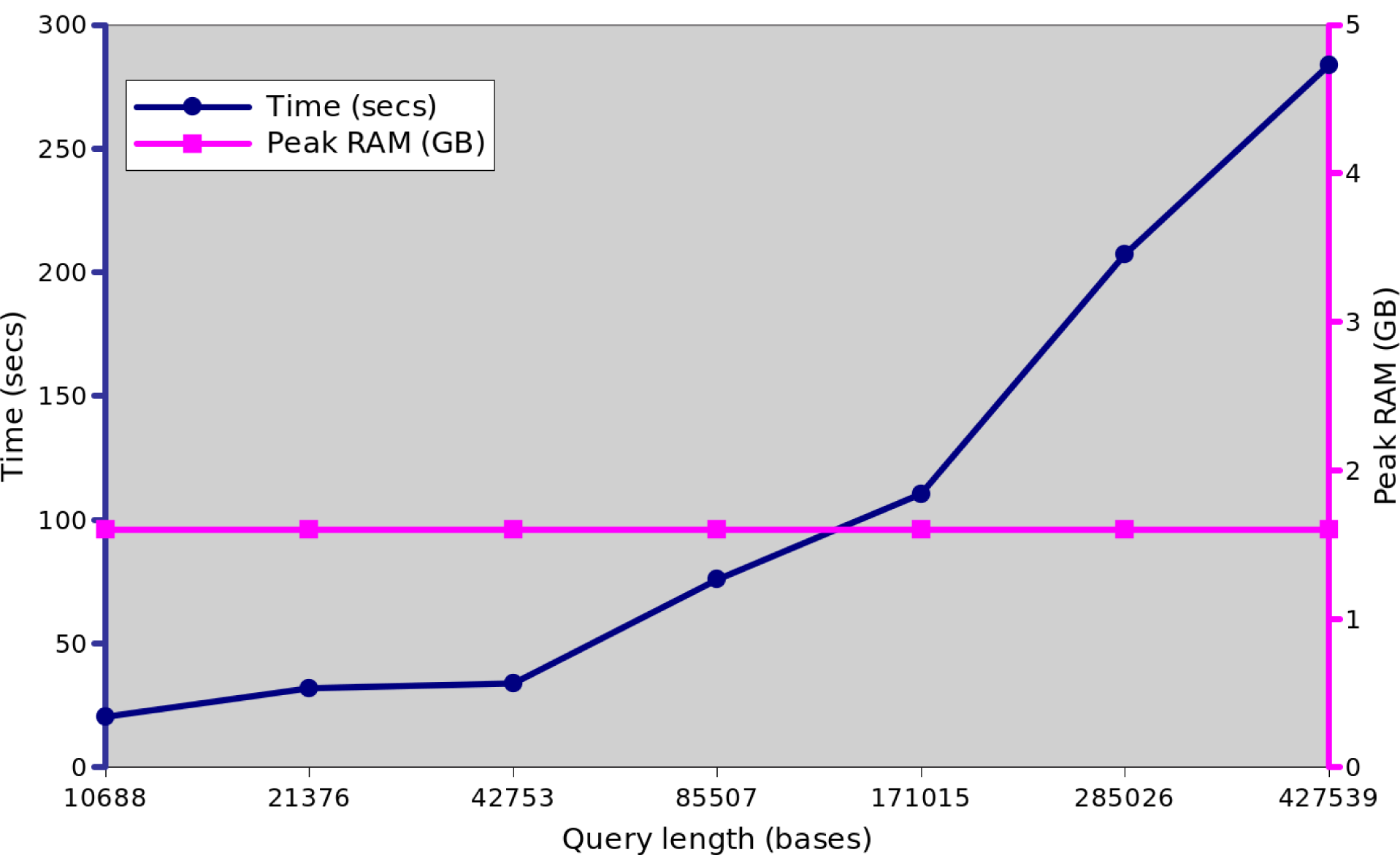
Time (seconds) and peak memory usage (GB) with increasing query length for “Sample sequence in the reference coordinate” (query #2) for 1000 Genomes chromosome 22 index. The time taken increases as the query length increases. But the memory usage remains constant regardless of the query length.

### The effect of number of variants on query time

We also evaluate how the number of variants in the position range affects the query time. For this benchmark, we performed the “All variants in the reference coordinate” query (query #6) because this query involves performing a breadth-first search in the graph to determine all variants in a region and the query time depends on the number of variants in the region. To perform queries on regions with different number of variants, we chose 1000 regions with a fixed size of the position range (≈ 42K bases) and start position chosen uniformly randomly across the chromosome.

Effect of the number of variants in the query region on query time is shown in Figure 4. The query time increases as the number of variants in the queried region increases. This is because when the number of variants is in a region is small the graph is sparser and faster to traverse and report all variants.

**Figure 4:**
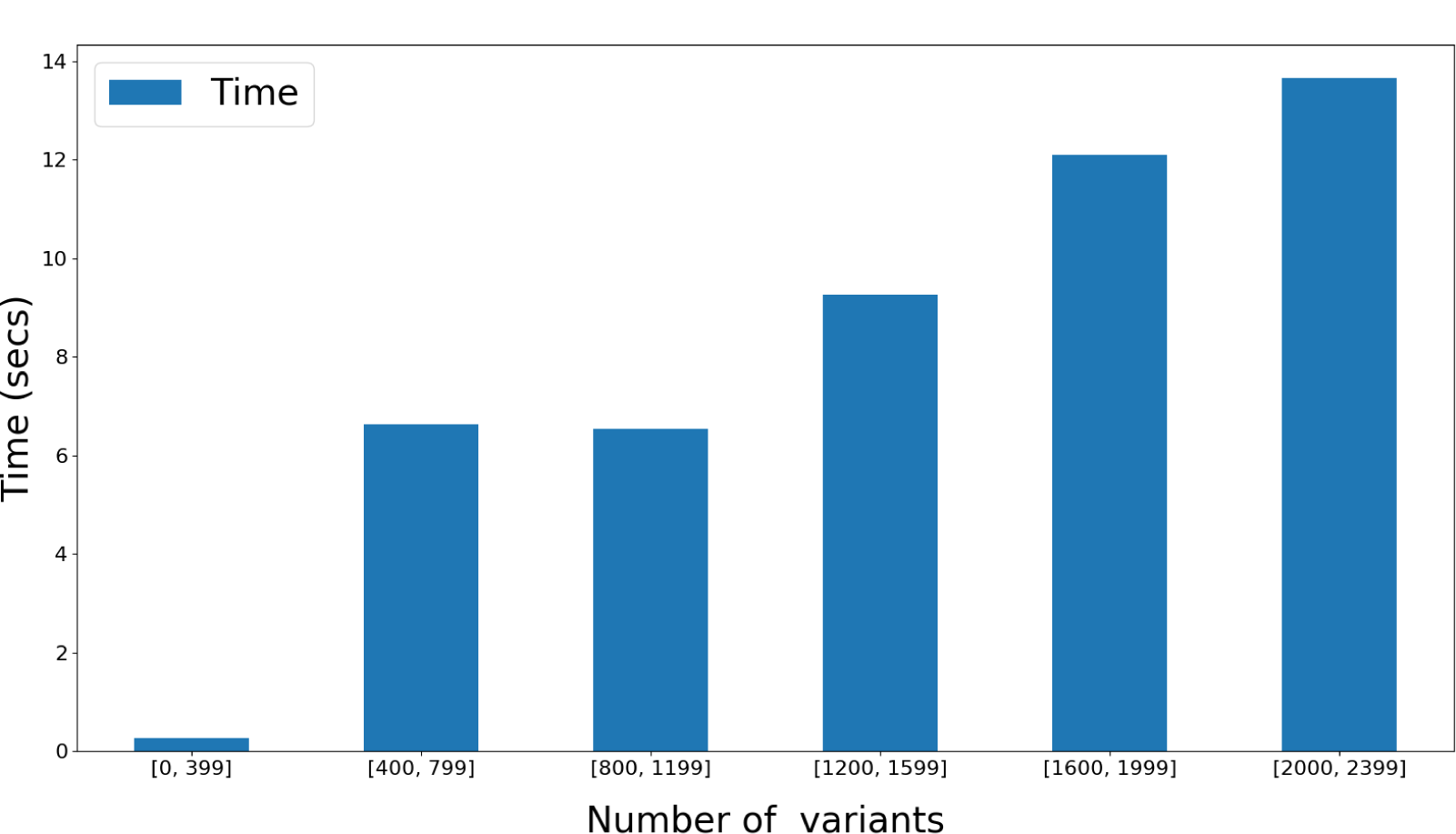
Time (seconds) and the number of variants in the query region for “All variants in the reference coordinate” (query #6) for 1000 Genomes chromosome 22 index. Query times are binned based on the number of variants in the query region and mean time is reported in each bin. The mean time increases as the number of variants increases in the region.

### Comparison with a reference-based variant index

To understand the overhead of maintaining thousands of coordinate systems on the performance of VariantStore, we compare VariantStore to a variant index that only indexes variants based on the reference coordinate system and supports a subset of variant queries. We use GTC [29] in our evaluation as it is the fastest and smallest reference-based variant index. We compare the query performance for two variant queries (Query#4 “Sample variants in ref coordinate” and Query#6 “All variants in ref coordinate”) supported by GTC.

GTC took 6× less time and an order of magnitude less space to construct and store the variant index compared to VariantStore. Furthermore, variant queries were also about an order of magnitude faster in GTC (see Figure 5). The slow performance of VariantStore is because of the overhead of maintaining the variation graph for representing multiple coordinate systems. During index creation, adding a variant requires splitting a reference node, adding the variant node in the graph, and updating the mapping function for all the samples corresponding to the variant. To query, we first map the variant position to a node in the graph using the position-index and then traverse the path in the graph to answer queries. On the other hand, adding and querying variants can be performed fairly efficiently using compressed bit vectors in reference-only variant indexes. If an application only requires querying variants based on a single reference coordinate system then GTC offers a space-efficient and faster alternative, but it does not support queries using per-sample coordinate systems or general genome graph traversals.

**Figure 5:**
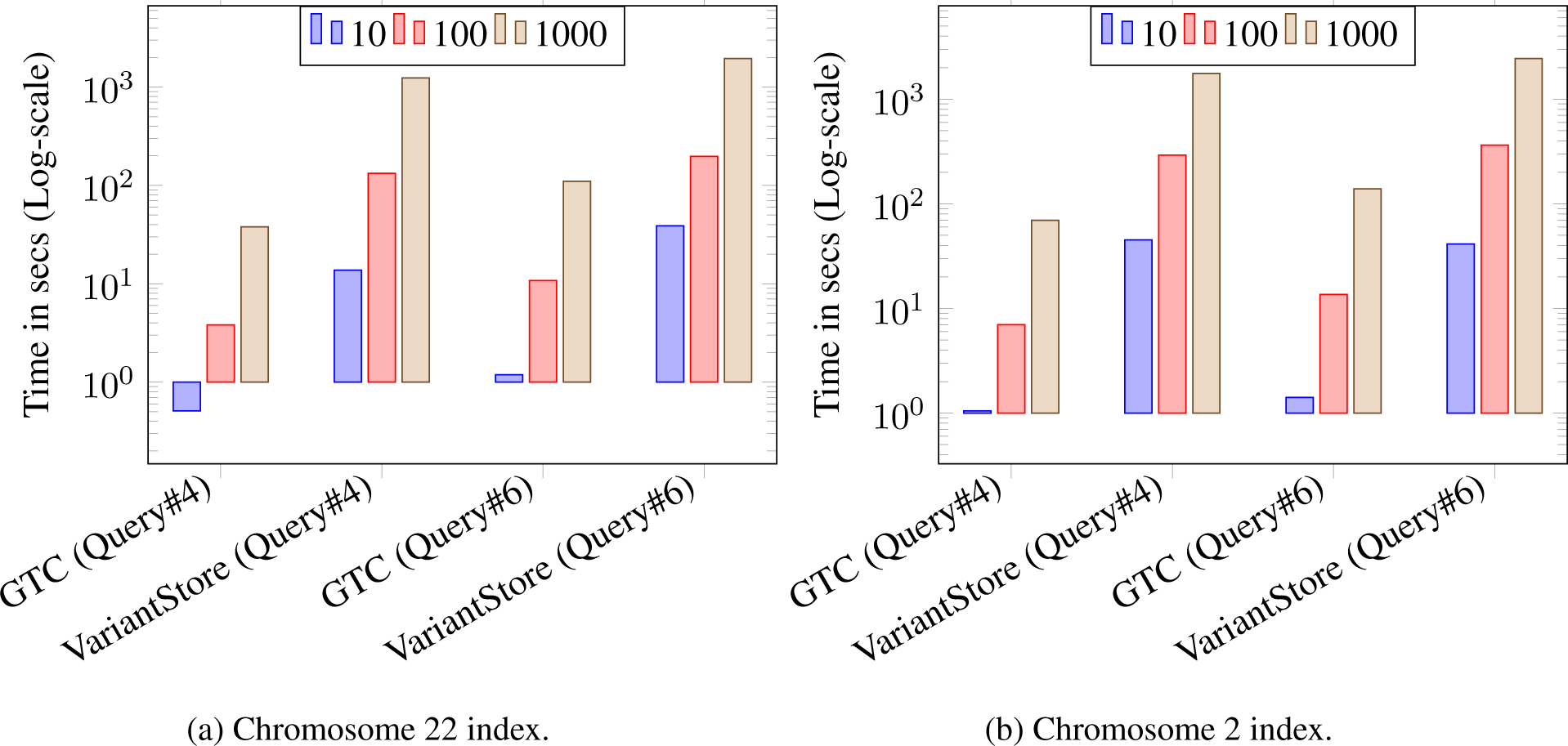
Time reported is the total time taken to execute 10, 100, and 1000 queries on 1000 genomes data. For all queries the query length is fixed to ≈ 42*K*. Query#4: Sample variants in ref coordinate and Query#6: All variants in ref coordinate

### Experimental hardware

All experiments on 1000 Genomes data were performed on an Intel Xeon CPU E5-2699A v4 @ 2.40GHz (44 cores and 56MB L3 cache) with 1TB RAM and a 7.3TB HGST HDN728080AL HDD running Ubuntu 18.04.2 LTS (Linux kernel 4.15.0-46-generic) and were run using a single thread. Benchmarks for TCGA data were performed on a cluster machine running AMD Opteron Processor 6220 @ 3GHz with 6MB L3 cache. TCGA data was stored on a remote disk and accessed via NFS.

## Discussion

We attribute the scalability and efficient index construction and query performance of VariantStore to the variant-oriented index design. VariantStore uses an inverted index from variants to the samples which scales efficiently when multiple samples share a variant which is often seen in genomic variation data. The inverted index design further allows us to build the position index only on the reference sequence and use graph traversal to transform the position in reference coordinates to sample coordinates.

All the supported variant queries look for variants in a contiguous region of the chromosome which allows VariantStore to partition the variation graph representation into small chunks based on the position of nodes in the chromosome and sequentially load only the relevant chunks into memory during a query. This makes VariantStore memory-efficient and scale to genomic variation data from hundreds of thousands of samples in future.

The variation graph representation in VariantStore is smaller and more efficient to construct than the representation in VG toolkit. It can be further used in read alignment as a replacement to the variation graph representation in the VG toolkit.

While the current implementation does not support adding new variants or updating the reference sequence in an existing VariantStore, this is not a fundamental limitation of the design. In future, we plan to extend the immutable version of VariantStore to support dynamic updates following the LSM-tree design [30].

## Methods

### Variation graph

A variation graph (VG) [14] (also defined as a genome graph in Kim et al. [17] and Rakocevic et al. [18]) is a directed, acyclic graph (DAG) *G* = (*N, E, P*) that embeds a set of DNA sequences. It comprises of a set of nodes *N*, a set of direct edges *E*, and a set of paths *P*. For DNA sequences, we use the alphabet {A, C, G, T, N}. Each *n*_*i*_ ∈ *N* represents a sequence *seq*(*n*_*i*_). Edges in the graph connect nodes that are followed on a path. Each sample in the VCF file follows a path through the variation graph. The embedded sequence given by the path is the sample sequence. Given that a variation graph is a directed graph, edges can be traversed in only one direction. Although, not applicable to VariantStore, an edge can also be traversed in the reverse direction when the variation graph is used for read alignment [14, 17, 18].

Each path *p* = *n*_*s*_*n*_1_ … *n*_*p*_*n*_*d*_ in the graph is an embedded sequence defined as a sequence of nodes between a source node *n*_*s*_ and a destination node *n*_*d*_. Nodes on a path are assigned positions based on the coordinate systems of sequences they represent [31]. The position of a node on a path is the sum of the lengths of the sequences represented by nodes traversed to get to the node on the path. For a path *p* = *n*_1_ … *n*_*p*_, position *P* (*n*_*p*_) is 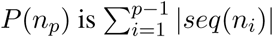. A node in the graph can appear in multiple paths and therefore can have multiple positions based on different coordinate systems.

An initial variation graph is constructed with a single node and no edges using a linear reference sequence and a reference genome coordinate system. Variants are added to the variation graph from one or more VCF files [13]. A variant is encoded by a node in the variation graph that represents the variant sequence and is connected to nodes representing the reference sequence via directed edges. Each variant in the VCF file splits an existing reference sequence node into two (or three in some cases) and joins them via an alternative path corresponding to the variant. For example, a substitution or deletion can cause an existing reference node to be split into three parts (Figure 6). An insertion can cause the reference node to be split into two parts (Figure 6).

**Figure 6:**
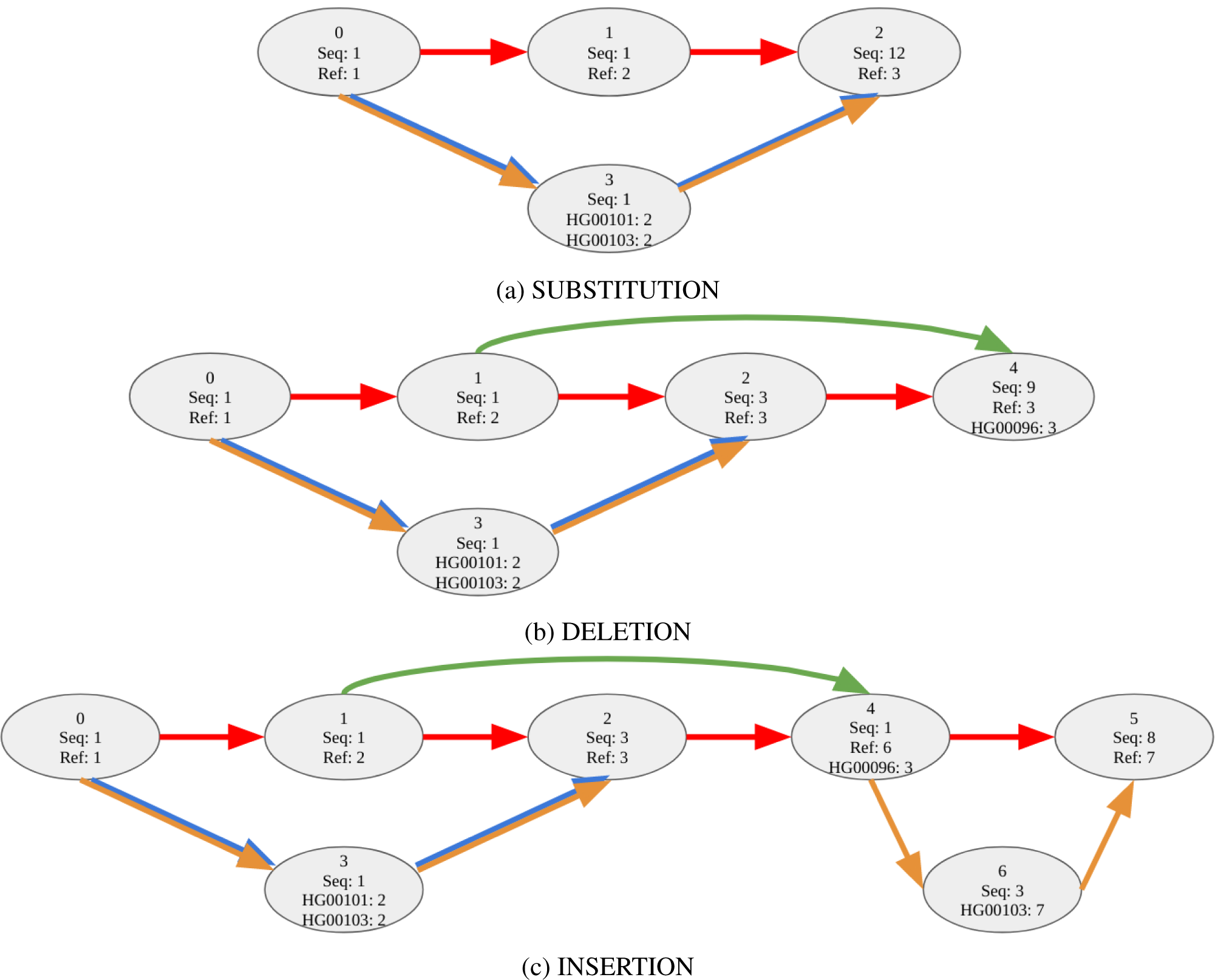
A variation graph with three input samples (HG00096, HG00101, HG00103) showing the encoding of substitutions, insertions, and deletions as stated in Table 2. Fig. (a) shows the substitution, (b) shows the deletion, and (c) shows the insertion. Edges are colored (or multi-colored) to show the path taken by reference and samples through the graph. (Ref: red, HG00096: green, HG00101: blue, HG00103: brown). Samples with no variant at a node follow the reference path, e.g., sample HG00096 will follow the reference path between nodes 0 *→* 1 and 4 *→* 5. Each node contains node id, the length of the sequence it represents, and a list of samples and their positions.

### Representing multiple coordinate systems in a variation graph

Representing multiple coordinates systems in a variation graph poses challenges that are not present in linear reference genomes. First, a node can appear on multiple paths at a different position on each path. Second, given that a variation graph can contain thousands of paths and coordinate systems it would be non-trivial to maintain a position index to quickly get to a node corresponding to a path and position.

**Table 2:** Variants ordered by the position in the reference genome for three samples (HG00096, HG00101, HG00103). Each variant has the list of samples that contain the variant.

| Reference sequence |  |  | CAATTTGCTGATCT |  |  |  |
| --- | --- | --- | --- | --- | --- | --- |
| Position | Reference seq. | Alternative seq. | HG00096 | HG00101 | HG00103 | Variant type |
| 2 | A | G | 0 | 1 | 1 | SUBSTITUTION |
| 2 | AATT | A | 1 | 0 | 0 | DELETION |
| 6 | T | TACG | 0 | 0 | 1 | INSERTION |

Much work has been done devising efficient approaches to handle multiple coordinate systems. Rand et al. [31] introduced the offset-based coordinate system. VG toolkit [26] implemented the multiple coordinate system by explicitly storing a list of node identifiers and node offsets for each path in the variation graph. They store the list of node identifiers as integer vectors using the succinct data structure library (SDSL [32]). We call this an *explicit-path representation*.

However, storing the list of nodes on each path explicitly can become a bottleneck as the number of input paths increases. Moreover, nodes that appear on multiple paths are stored multiple times causing redundancy in storage. For a set of *N* variants and *S* samples the space required to store the explicit-path representation is *O*(*SN*) since each variant creates a constant number of new nodes in the variation graph.

### VariantStore

We describe how we represent a variation graph in VariantStore and maintain multiple coordinate systems efficiently. We then describe how we build a position index using succinct data structures.

The variation graph representation is divided into three components:

1. Variation graph topology
2. Sequence buffer
3. List of variation graph nodes

#### Variation graph topology

A variation graph constructed by inserting variants from VCF files often shows high sparsity (the number of edges is close to the number of nodes). For example, the ratio of the number of edges to nodes in the variation graph on 1000 Genomes data [27] is close to 1. Given the sparsity of the graph, we store the topology of the variation graph in a representation optimized for sparse graphs.

Our graph representation uses the counting quotient filter (CQF) [33] as the underlying container for storing nodes and their outgoing neighbors. The CQF is a compact representation of a multiset *S*. Thus a CQF supports inserts and queries. A query for an item *x* returns the number of instances of *x* in *S*. The CQF uses a variable-length encoding scheme to count the multiplicity of items. In our graph representation, we encode the node as the item and id of the outgoing neighbor as the count of the item in the CQF.

For nodes with a single outgoing neighbor, we store the node and its neighbor together (without an indirection) so we can access them quickly. For nodes with more than one outgoing neighbor we store outgoing neighbors in separates lists. In the variation graph, most nodes have a single outgoing neighbor and using this compact and optimized representation we achieve cache efficient and fast traversal of the graph.

We use the version of the counting quotient filter with no false-positives to map a node to its outgoing edge(s). We store the node id as the key in the CQF and if there is only one outgoing edge we encode the outgoing neighbor id as the count of the key. If there are more than one outgoing edges we use indirection. We maintain a list of vectors where each vector contains a list of outgoing neighbor identifiers corresponding to a node. We store the node id as the key and the offset in the list of vectors (or index of the vector containing the list of outgoing neighbor ids) as the count of the key.

#### Sequence buffer

The sequence buffer contains the reference sequence and all variant sequences corresponding to each substitution and insertion variant. All sequences are encoded using 3-bit characters in an integer vector from SDSL library [32, 34]. The integer vector initially only contains the reference sequence. Sequences from incoming variants are appended to the integer vector. Once all variants are inserted the integer vector is bit compressed before being written to disk.

#### List of variation graph nodes

Each node in the variation graph contains an offset and length. The offset points to the start of the sequence in the sequence buffer and length is the number of nucleotides in the sequence starting from the offset. This uniquely identifies a node sequence in the sequence buffer.

At each node we also store a list of sample identifiers that have the variant, position of the node on all those sample paths, and phasing information from the VCF file corresponding to each sample.

Our representation of the list of samples is based on two observations. First, multiple samples share a variant and storing a list of sample identifiers for each variant is space inefficient. Instead, we store a bit vector of length equal to the number of samples and set bits corresponding to the present samples in the bit vector. Second, multiple variants share the same set of samples. We define an equivalence relation ∼ over the set of variants. Let *E*(*v*) denote the function that maps each variant to the set of samples that have the variant. We say that two variants are equivalent (i.e., *v*_1_ ∼ *v*_2_) if and only if *E*(*v*_1_) = *E*(*v*_2_). We refer to the set of samples shared by variants as the *sample class*. A unique id is assigned to each sample class and nodes store the sample class id instead of the whole sample class. This scheme has been employed previously by other colored de Bruijn graph representation tools [35–38] for efficiently maintaining a mapping from *k*-mers (a *k*-length substring sequence) to the set of samples where *k*-mers appear.

Phasing information is encoded using 3 bits. Position and phasing information corresponding to each sample in the list of samples is stored as tuples. Tuples are stored in the same order as the samples appear in the sample class bit vector. To retrieve the tuple corresponding to a sample a rank operation is performed on the sample bit vector to determine the rank of the sample. Using the rank output, a select operation is performed on the tuple list to determine the tuple corresponding to a sample.

Variation graph nodes are stored as protocol buffer objects using Google’s open-source protocol buffers library. Every time a new node is created we instantiate a new protocol buffer object in memory. We compress the protocol buffers before writing them to disk and decompress them while reading them back in memory.

For a set of *N* variants and *S* samples where each variant is shared by *P* samples on average, each node contains information about *P* samples and storing *O*(*N*) nodes (a constant number of nodes for each variant) the space required to store the variation graph representation in VariantStore is *O*(*NP*). When *P* = 1 (i.e., no two samples share a variant) the space required by the variation graph representation becomes *O*(*N*).

#### Position index

In order to answer variant queries we need an index to quickly locate nodes in the graph corresponding to input positions. These positions can be specified in multiple coordinate systems, i.e., in the coordinate system of the reference or a sample.

One way to index the variation graph is to store an ordered mapping from position to node identifier. We can perform a binary search in the map to find the position closest to the queried position and the corresponding node id in the graph. However, given that there are multiple coordinate systems in the variation graph we can not create a single mapping with a global ordering. Keeping a separate position index for each coordinate system will require space equal to the explicit-path representation.

In VariantStore, we maintain a mapping of positions to node identifiers only for the reference coordinate system. All nodes on the reference sequence path are present in the mapping. If the queried position is in the reference coordinate system we use the mapping to locate the node in the graph. However, if the queried position is in a sample’s coordinate system, we first locate the node in the graph corresponding to the same position in the reference coordinate system. Then we perform a local search by traversing the sample path from that node to determine the node corresponding to the position in sample’s coordinate system. The local graph search incurs a small, one-time cost because sample nodes are rarely far from a reference node and is amortized against future searches in the sample’s coordinate system.

We create the position index using a bit vector called the position-bv of length equal to the reference sequence length and a list of node identifiers on the reference path in the increasing order by their reference sequence positions. For every node in the list we set the bit corresponding to the node’s position in the position-bv. There is a one-to-one correspondence between every set bit in the position-bv and node positions in the list. We store the position-bv using a bit vector and node list as an integer vector from the SDSL library [32, 34].

### Variation graph construction

We construct the variation graph by inserting variants from a VCF file. Each variant has a position in the reference genome, alternative sequence (except in case of a deletion), and a list of samples with phasing information for each sample.

Based on the position of the variant we split an existing reference node that contains the sequence at that position in the graph. We update the split nodes on the reference path with new sequence buffer offsets, lengths, and node positions (based on the reference coordinate system). We then append the alternative sequence to the sequence buffer and create an alternative node with the offset and length of the alternative sequence. We then add the list of tuples (position, phasing info) for each sample.

We also need to determine the position of the alternative node on the path of each sample that contains the variant. One way to determine the position of the node for each sample would be to backtrack in the graph to determine a previous node that contains a sample variant and the absolute position of that node in the sample’s coordinate system. If no node is found with a sample variant we trace all the way back to the source of the graph. We would then traverse the sample path forward up to the new alternative node and compute the position. This backtracking process would need to be performed once for each sample that contains the variant. This would slow down adding a new variant and cause the construction process to not scale well with increasing number of samples.

Instead, we construct the variation graph in two phases to avoid the backtracking process. In the first phase, while adding variants we do not update the position of nodes on sample paths. We only maintain the position of nodes on the reference path because that does not require backtracking. In the second phase, we perform a breadth-first traversal of the variation graph starting from the source node and update the position of nodes on sample paths.

During the breadth-first traversal we maintain a delta value for each sample in the VCF file. At any node, the *delta value* is the difference between the position of the node in the reference coordinate and the sample’s coordinate. During the traversal, we update sample positions for each node based on the current delta value and reference coordinate value. Algorithm 1 gives the pseudocode of the algorithm.

#### Algorithm 1

Pseudocode to fix sample positions in the variation graph. A node corresponding to a variant contains the list of sample identifiers that have the variant and their respective positions in sample paths. A node corresponding to the reference sequence contains the position in the reference path and optionally a list of sample identifiers if it also represents a delete variant.

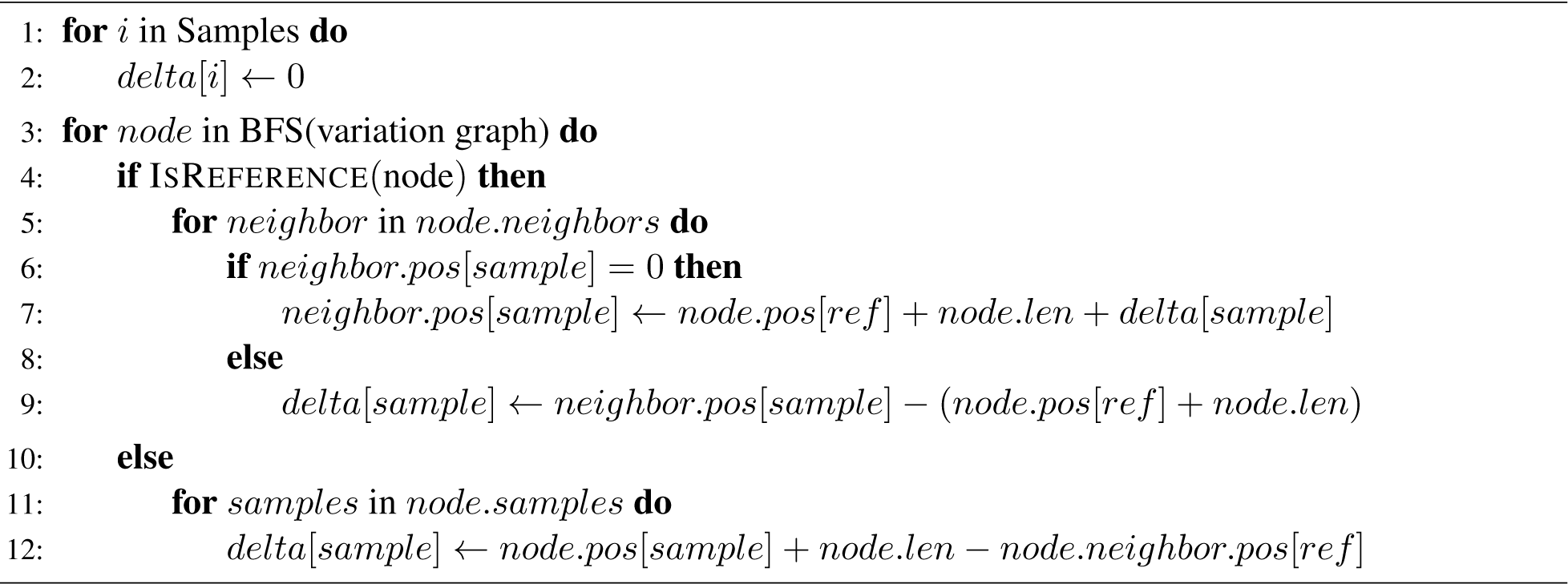

### Position index construction

In the position index, we maintain a mapping from positions of nodes on the reference path to corresponding node identifiers in the graph. Node positions are stored in a “position-bv” bit vector of size equal to the length of the reference sequence and node identifies are stored in a list. To construct the position index, we follow the reference path starting from the source node in the graph and for every node on the path we set the corresponding position bit in the position-bv and add the node identifier to the list. Node identifiers are stored in the order of their position on the reference path.

### Variant queries

A query is performed in two steps. We first perform a predecessor search (largest item smaller than or equal to the queried item) using the queried position in the position index to locate the node *n*_*p*_ with the highest position smaller than or equal to the queried position *pos*. The predecessor search is implemented using the rank operation on position-bv. For bit vector *B*[0, …, *n*], *RANK*(*j*) returns the number of 1s in prefix *B*[0, …, *j*] of *B*. An RRR compressed position-bv supports rank operation in constant time [32, 39]. The rank of *pos* in position-bv corresponds to the index of the node id in the node list. Figure 7a shows a sample query in the position index.

**Figure 7:**
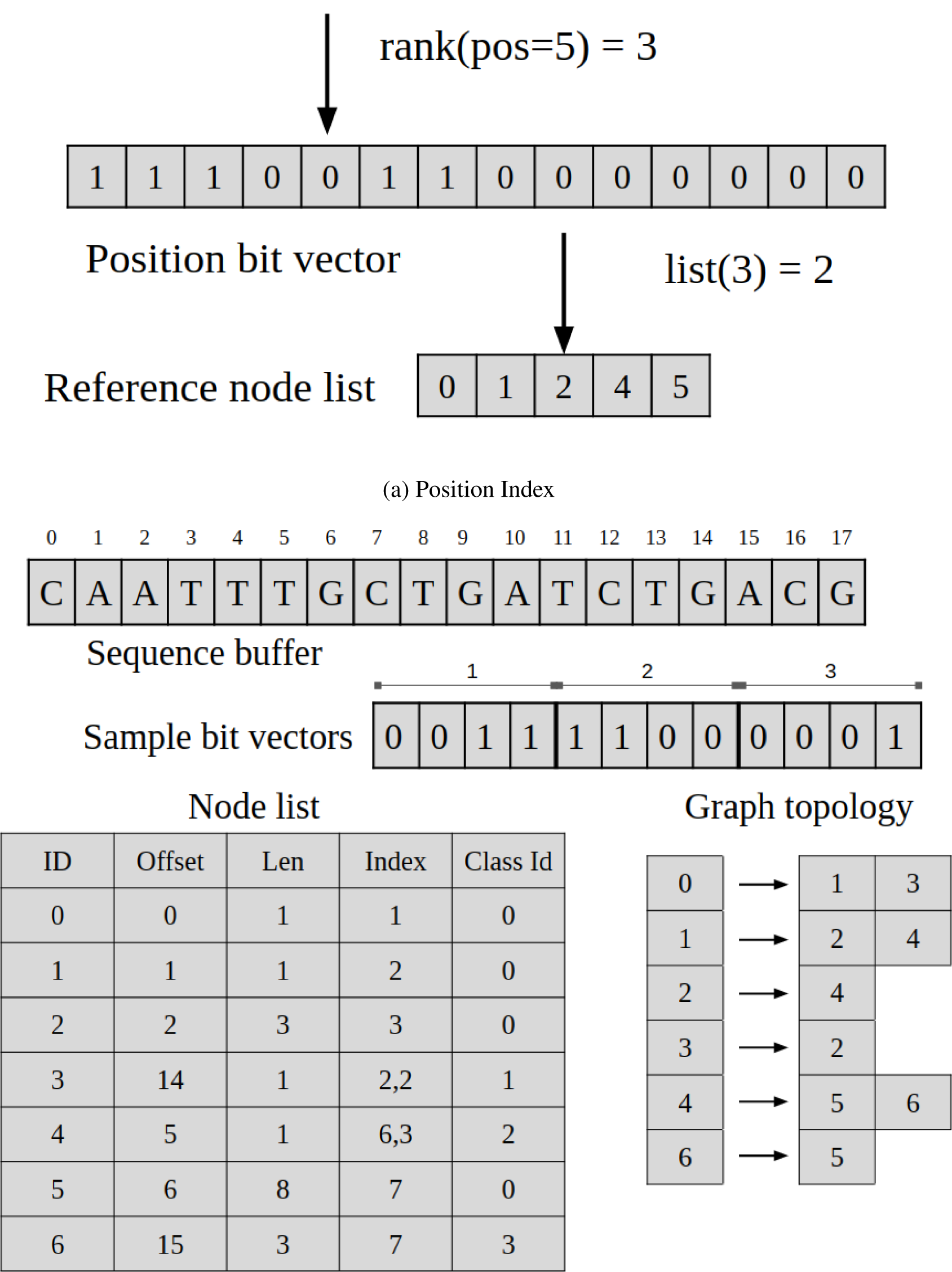
Position index and variation graph representation in VariantStore for the sample graph from Figure 6. (a) Shows the query operation for finding the node at position 5 in the sequence. (b) Phasing information is omitted from the node list for simplicity of the figure. In implementation, phasing information using three bits for each sample in each node.

Based on how reference nodes are split while adding variants the sequence starting at *pos* will be contained in the node *n*_*p*_. All queries are then answered by traversing the graph either by following a specific path (reference or a sample) or a breadth-first traversal and filtering nodes based on query options.

If the queried position is based on the reference coordinate system then we can directly use *n*_*p*_ as the start node for graph traversal. However, if the position is based on a sample coordinate then we perform a local search in the graph starting from *n*_*p*_ to determine the start node based on the sample coordinate.

### Memory-efficient construction and query

In the variation graph representation, the biggest component in terms of space is the list of variation graph nodes stored as Google protobuf objects. These node objects contain the sequence information and the list of sample positions and phasing information. For 1000 Genomes data, the space required for variation graph nodes is ≈ 87% to 92% of the total space in VariantStore. However, keeping the full list of node objects in memory during construction or query is not necessary and would make these processes memory inefficient.

To perform memory efficient construction and query, we store and serialize these nodes in small chunks usually containing ≈ 200K nodes (the number of nodes in a chunk varies based on the data to keep the size to a few MBs). Nodes in and across these chunks are kept in their creation order (which is roughly the breadth-first traversal order). Therefore, during a breadth-first traversal of the graph we only need to load these chunk in sequential order.

During construction, we only keep two chunks in memory, the current active chunk and the previous one. All chunks before the previous chunk are written to disk. In the second phase of the construction when we update sample positions and during the position index creation we perform a breadth-first traversal on the graph and load chunks in sequential order.

Variant queries involve traversing a path in the graph between a start and an end position or exploring the graph locally around a start position. All these queries require bounded exploration of the graph for which we only need to look into one or a few chunks.

To perform queries with a constant memory we only load the position index and variation graph topology in memory and keep the node chunks on disk. We use the index and the graph topology to determine the set of nodes to look at to answer the query. We then load appropriate chunks from disk which contain the start and end nodes in the query range. For queries involving local exploration of the graph we load the chunk containing the start node. During the exploration, we load new chunks lazily as needed. At any time during the query, we only maintain two contiguous chunks in memory.

## Data availability

Code and data are available at https://github.com/Kingsford-Group/variantstore

## Acknowledgements

The results published here are in whole or part based upon data generated by The Cancer Genome Atlas (dbGaP accession phs000178) managed by the NCI and NHGRI. Information about TCGA can be found at http://cancergenome.nih.gov. The 1000 Genomes data used for the analyses described in this manuscript were obtained from https://www.internationalgenome.org/. This research is funded in part by the Gordon and Betty Moore Foundation’s Data-Driven Discovery Initiative through Grant GBMF4554 to C.K. and by the US National Institutes of Health (R01GM122935). We would also like to thank Shawn Baker for many helpful discussions, and Guillaume Marçais and Yutong Qiu for comments on the manuscript.

